# S-IRFindeR: stable and accurate measurement of intron retention

**DOI:** 10.1101/2020.06.25.164699

**Authors:** Lucile Broseus, William Ritchie

## Abstract

Accurate quantification of intron retention levels is currently the crux for detecting and interpreting the function of retained introns. Using both simulated and real RNA-seq datasets, we show that current methods suffer from several biases and artefacts, which impair the analysis of intron retention. We designed a new approach to measure intron retention levels called the Stable Intron Retention ratio that we have implemented in a novel algorithm to detect and measure intron retention called S-IRFindeR. We demonstrate that it provides a significant improvement in accuracy, higher consistency between replicates and agreement with IR-levels computed from long-read sequencing data.

S-IRFindeR is freely available at: https://github.com/lbroseus/SIRFindeR/.

## Background

Intron Retention (IR) is a type of alternative splicing that is gaining increased interest in human health and disease research. Originally described in plants and viruses, IR has now been shown to be a common form of alternative splicing in mammalian systems [1,2]. However, quantifying IR levels poses several specific difficulties [2–4]. Introns are highly heterogeneous genomic regions, both in length and sequence features. In mammals, IR levels are generally low and thereby subject to incomplete coverage and higher count overdispersion [5,6].

Two approaches have been proposed to address the specifics of IR-level quantitation [4]: an intronic-tuned version of the Percentage Spliced-In (PSI) value [7,8], and our own IRratio [2] **(Table 1**). Both methods estimate the portion of transcripts including the intron (*intronic abundance*) and the portion of spliced transcripts (*splicing abundance*), and compute the retention fraction as the ratio: intronic abundance / (splicing abundance + intronic abundance), but they differ markedly in their strategy to measure these two quantities (see [4] for a review).

**Table 1:**
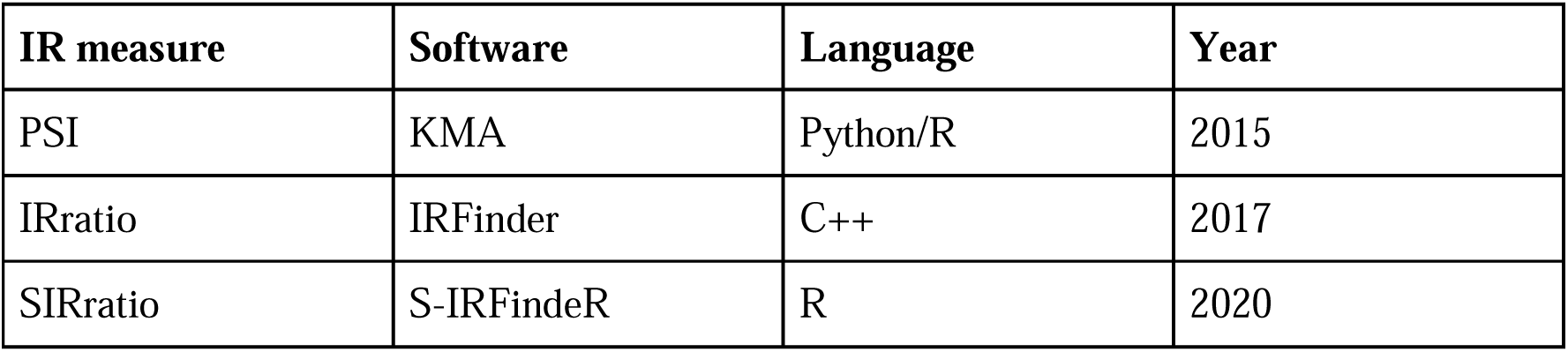
Existing measures to estimate IR levels from short RNA-seq data and their implementation.

Though these methods have been recently used to quantify IR events in a wide range of studies [9–16], their capacity to reflect true IR transcripts has not yet undergone thorough validation [3,17]. In fact, several authors including ourselves highlight the high-variability and lack of reproducibility of IR level estimates [3,18].

## Results and Discussion

By analysing both simulated and real data (Supplementary Methods), we found that current estimators frequently report aberrant IR values. Many of these are known to be caused by splicing events that are absent from reference annotations [4] (especially novel 3’ or 5’ donor sites). In addition, we found that the IRratio and PSI calculated by KMA provided poor estimates of normal splicing and intronic abundance and produced aberrant zero and one values which led to inconsistent IR levels (**Figure 1A** and **Supplementary Figures 1 and 2**). These extreme values of “1” and “0” are a major handicap for downstream analyses. An artifactual value of 1 can be falsely interpreted as “all transcripts retain this intron” and will thus be considered as prime candidates for further bioinformatics analysis or even experimental validation. A false value of 0 will constitute a false negative but in addition will perturb comparison of IR between samples. We also found that both the IRratio and PSI had a tendency to underestimate the level of the longest introns (**Figure 1B**). More generally, inconsistent estimates can affect the interpretation of IR events and the identification of relevant molecular signatures (**Figures 2A and 2B** and **Supplementary Figure 8**).

**Figure 1:**
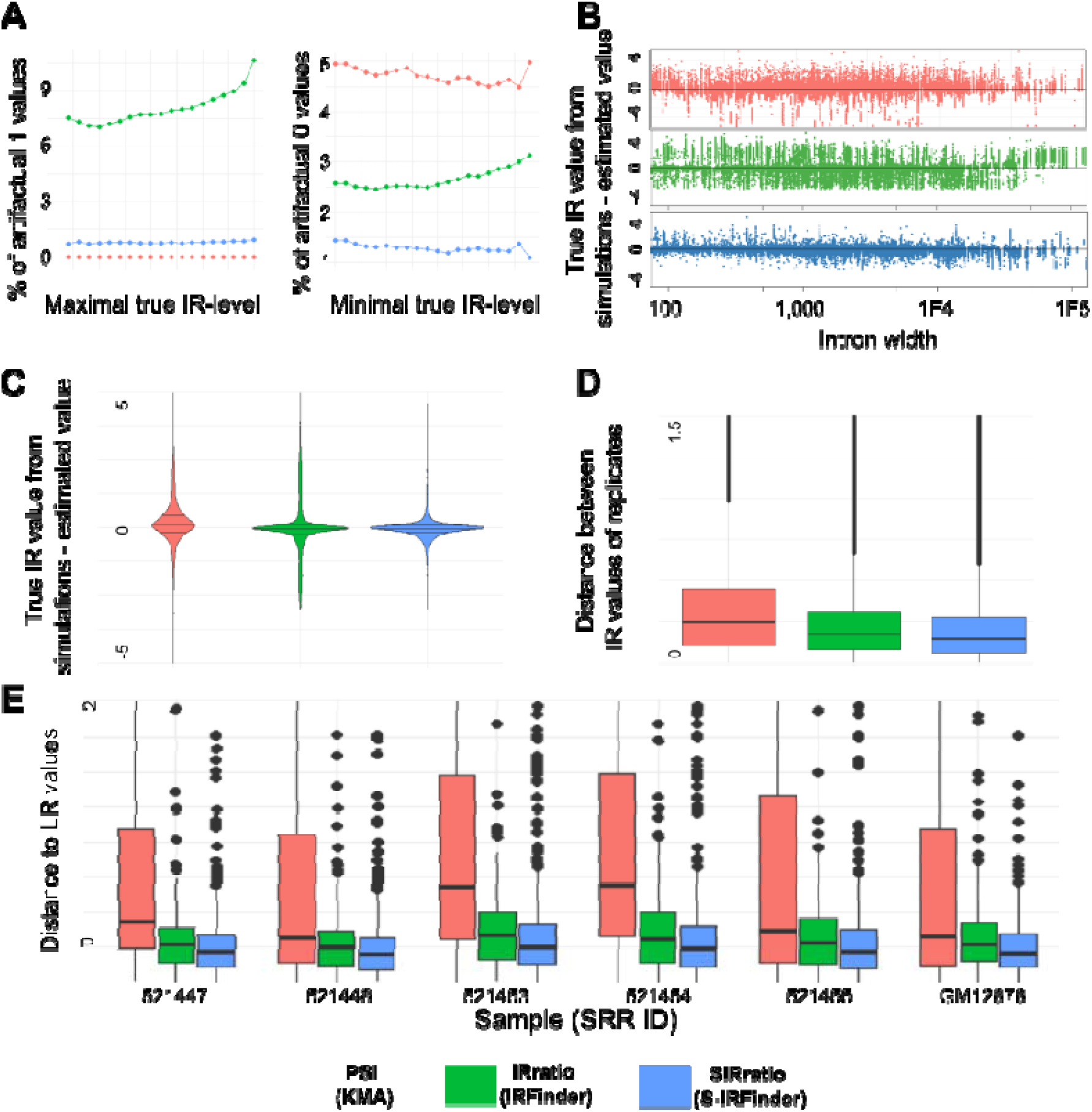
Comparison of the three IR-measures on real and simulated RNA-seq data. **A.** Percentage of artifactual “0” and “1” IR values. **B.** The effect of intron length on the difference between the true IR-levels and their estimates. Both the PSI and the IRratio tend to underestimate the retention level of the longest introns. **C.** Distribution of differences between the true IR-levels and their estimates on a simulated RNA-seq experiment. **D.** Real data: overall distribution of distances between five technical replicates from the GM12878 cell line. **E**. Real data: distribution of distances between IR-levels obtained from Oxford Nanopore long read data and those computed from short read data on the same GM12878 human cell line. Additional results and figures can be found in Supplementary Materials.

**Figure 2.**
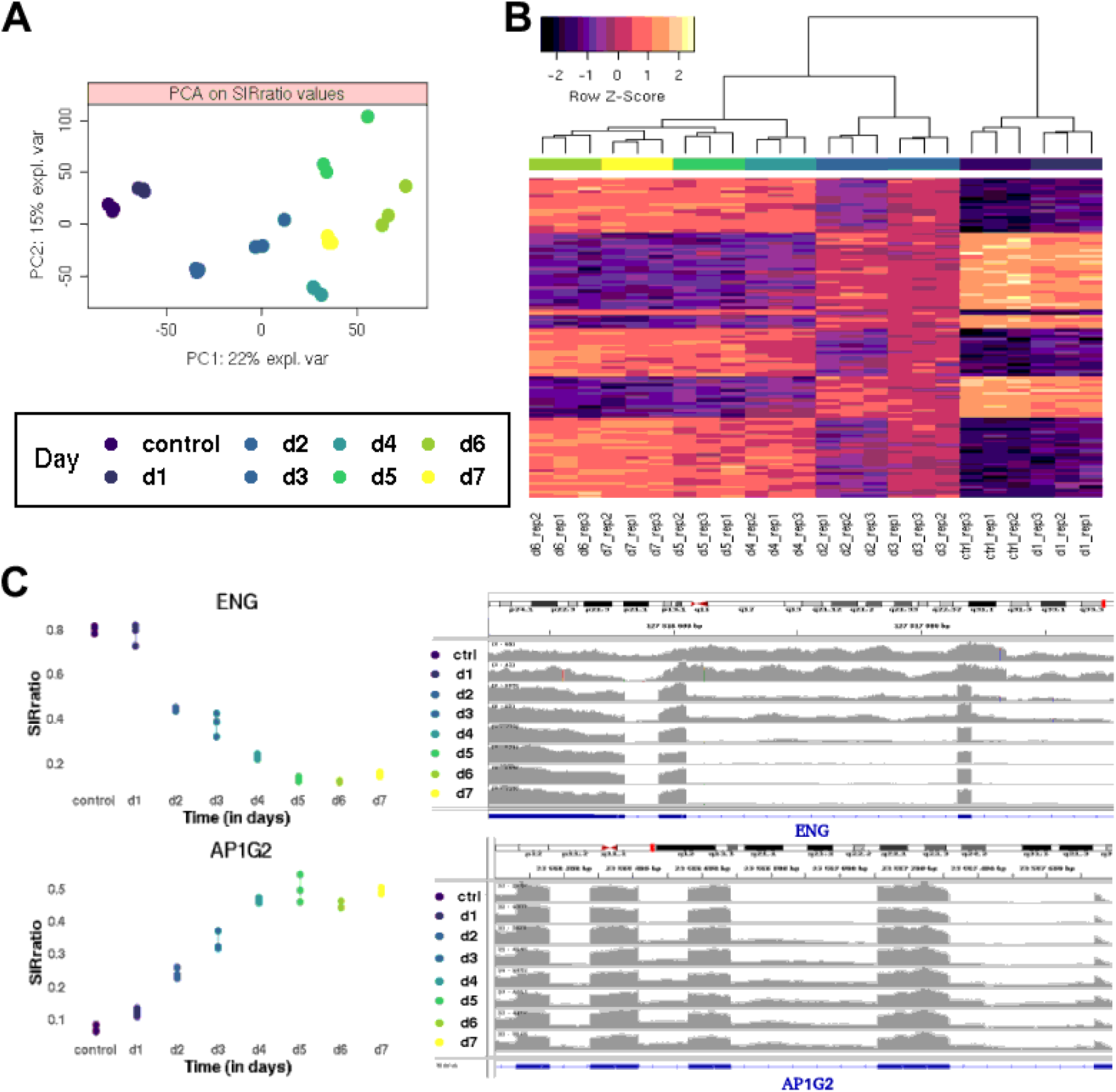
Temporal variations of IR-levels during the EMT induced in a Human non-small cell lung cancer cell line H358. **A.** Principal component analysis of SIRratio values across time points. The first component captures the temporal nature of the data, and the progress of the EMT. Values obtained with the IRratio are noisier and harder to interpret (cf: Supplementary Figure 8). **B.** Heatmap of SIRratio values and hierarchical clustering on the 100 most influential retained introns on the first component, selected by sparse PCA [36,37]. **C.** Illustration of the evolution of coverage profiles and SIRratio levels on two of the introns driving the first component of the sparse PCA: Intron 13 from gene Endoglin (ENG, Chr9: 127,815,012-127,854,756, reverse strand) and Intron 2 from gene AP1G2 (Chr14: 23,559,565-23,568,070 reverse strand).

Taking into consideration these biases, we shaped a novel estimator, coined SIRratio for Stable Intron Retention ratio. We used a shrinkage approach to share coverage information at the gene level and improve the stability and interpretability of normal splicing estimates. We integrated this novel metric into a new version of IRFinder that we called S-IRFinder. To assess the efficiency of the SIRatio, we first generated multiple sets of simulated IR events with varying coverage and retention levels (**Supplementary Materials**). We found that the IR levels found by the SIRatio were closer to the real levels (**Figure 1C and Supplementary Figure 2**). We also found that the SIRatio was more consistent between technical replicates than the IRratio or the PSI (**Figure 1D, Supplementary Figures 3 & 4**).

Finally, for further validation on real data, we made use of direct RNA sequencing, developed by Oxford Nanopore Technologies. This technology represents a unique opportunity for the detection, validation and quantification of IR because it can capture entire IR transcripts in one sequencing read. S-IRFinder now includes a pipeline and guidelines for detecting IR events in long read data (Supplementary Material and on the S-IRFinder repository). We took advantage of matched short and long read datasets to confirm that the SIRratio provides sensible results on real data as well (**Figure 1E**). The S-IRFinder estimates were systematically closer to the long read counts than the other two estimates.

Numerous reports have demonstrated a regulatory role for IR in cell differentiation [9,19,20] and cancer [21–23,16]. However contrasting the IR levels of different stages of differentiation or conditions is currently a weak point of all IR detection and estimation algorithms [4]. To exemplify the practical value of the SIRratio, we took advantage of a large public time course experiment to investigate IR levels during the Epithelial-to-Mesenchymal (EMT) transition. Using principal component and clustering analyses on SIRratios, we observed that the SIRratio was able to supply immediately intelligible and biologically meaningful results. These straightforward analyses highlighted clear temporal and stage patterns (cf: **Figure 2A and 2B**), which were missed by the IRratio (cf: **Supplementary Figure 8**).

## Conclusion

Until recently, IR detection ran parallel with the analysis of other splicing events without taking into account inherent difficulties in measuring intronic expression. As a result, IR had been systematically underestimated. Despite the recent development of specialized software for detecting IR, the measurement of IR levels has been problematic. Poor estimates of IR levels may distort the downstream analysis of enriched gene families, confound differential analysis of IR between biological conditions and lead to erroneous conclusions about the impact of IR in normal biology and disease. We developed a new metric, the SIRratio that stabilizes the estimates of IR by using a localized shrinkage approach and a more careful definition of intronic regions. This approach gives more accurate IR estimates when compared with simulated IR transcripts but also with third-generation long-read technology. It also gives more stable IR estimates, which facilitates the implementation of transcriptome-wide IR screenings and search for relevant biomarkers. We therefore believe that these improvements can favor routine investigation of IR events and help assess the role of intron retention in biology and disease.

Along with the capability to use long-read sequencing technologies to detect and measure IR, the SIRratio is now implemented in S-IRFindeR that we recommend as an alternative to our own IRFinder algorithm.

## Methods

Building on IRFinder’s approach [2,4] and taking into account several biases which can adversely affect current IR measures, we devised a new approach for quantifying IR-levels, called S-IRFindeR.

### Data-driven refinement of intron annotation

Default in intron annotation is a major cause for inaccurate IR values [17]. Current methods address this issue by detecting spurious coverage patterns (eg: read coverage entropy [24], probabilistic test [7]) that may impair quantitation, and exclude affected introns from downstream analyses. To allow considering even introns with unexpected alternative splicing events, we propose instead a procedure to polish intron intervals using sample-specific junctions (cf: **Supplementary Materials**).

### Stabilized estimation of IR-levels

Observing that splice junction counts and the median intron depth could provide unstable measures of the abundances of spliced and IR transcripts respectively, we stepped on popular shrinkage [25,26] and resampling techniques [27] to formulate a novel metric, the SIRratio (cf: **Supplementary Materials**).

### Benchmark

To evaluate the ability of the three IR-measures to reflect the true proportion of IR-transcripts, we generated 45 RNA-seq samples with known IR-levels, alternative exon splicing and varying gene coverage (cf: **Supplementary Materials**).

We then sought to extend our study to real RNA-seq data. Assuming IR-levels should be similar across technical replicates, we made use of five RNA-seq Poly-A runs, with varying library size, available from the human cell line GM12878 [28] to evaluate the capacity of each method to provide reproducible results. In order to evaluate reproducibility, we computed the distances between technical replicates [29] (**cf: Figure 1C, Supplementary Figure 3 and Supplementary Table 2**) in addition to correlation coefficients (**cf: Supplementary Figure 4**). What also motivated the choice for this RNA-seq experiment, was the availability of a deep long read experiment (over 10 million long reads) from the same cell line [30]. Oxford Nanopore data indeed allow sensible quantitation of RNA levels [31,32], demonstrate high correlation with RT-qPCR measurements [33] and offer a wider and unbiased panel of intron targets for validation. We designed and implemented a straightforward procedure to estimate intron retention rates from long reads (cf: **Validation using real Oxford Nanopore Long Read** in **Supplementary Materials).** As a means to ascertain short read estimates on real data, we then compared them to those computed from long-read data (**cf: Figure 1E and Supplementary Table 3**).

One of the assets of RNA-seq technologies for IR studies is to allow transcriptome-wide search for molecular signatures [34,12,16] and biomarkers [11]. To further assess the potential of the SIRratio and the benefits of accurate IR quantitation in large-scale studies, we applied our method to a large time course RNA-seq experiment carried on on H358 cells enduring Epithelial-to-Mesenchymal transition (EMT) [35]. We appraised the interpretability of the SIRratio and IRratio estimates using straightforward principal component analysis (PCA) and hierarchical clustering on SIRratios selected by a sparse principal component analysis [36,37] (cf: **Figure 2A and 2B** and **Supplementary Figure 8**). Additionally, this exploration allowed us to double-check that our estimator provides realistic and consistent values in real data settings (cf: **Figure 2C**).

