## Supplementary Material for "S-IRFindeR: stable and accurate measurement of intron retention"

**Evaluated IR-measures**

**Intronic PSI** values were computed using KMA [[1]](https://www.zotero.org/google-docs/?Ahd9oV) (version 0.1.0). We followed the instructions given in the vignette accompanying the package: reads were aligned using Bowtie2 [[2]](https://www.zotero.org/google-docs/?s1CMq7) (version 2.3.2) on an augmented transcriptome, and transcript-level quantification was performed with eXpress [[3]](https://www.zotero.org/google-docs/?joe5sR) (version 1.5.1).

**IRratio** values were obtained from IRFinder [[4]](https://www.zotero.org/google-docs/?SnIdEh) (version 1.2.4). We followed the steps described in the documentation (<https://github.com/williamritchie>).

**SIRratio** values were calculated with the help of the R package IRFindeR-S (version 0.1.0) using the exact same STAR alignments as obtained with IRFinder’s default parameters.

It is worth noting that **IntEREst** [[5]](https://www.zotero.org/google-docs/?RPa7QR) and **iRead** [[6]](https://www.zotero.org/google-docs/?8hWgew) do not estimate (relative) IR-levels, but rather compute normalised FPKM values, used for detecting IR events.

**Analysis of relative data**. As they are relative measures, constrained between 0 and 1, ratios should be transformed before computing the usual statistics (eg: arithmetic mean, median, standard deviations, euclidean distances) [[7,8]](https://www.zotero.org/google-docs/?Jdm5bm). Accordingly, all through the article, the Isometric Log-Ratio transformation (ilr) has been applied to all relative IR values. In our univariate case, it simply amounts to a rescaled logit transformation of each IR rate:

ilr(rate) = sqrt(0.5) x log($\frac{rate}{1-rate}$).

Zero values were set to 0.001, one values were set to 0.999.

**Additional Results**

**Supplementary Figure 1: Scatterplot of the true IR-levels against their estimates (Simulated data).** For each method, the global Spearman rank correlation coefficient (cor) is specified. In the middle plot, we observe that the IRratio gives many incorrect zero values (along the x-axis).

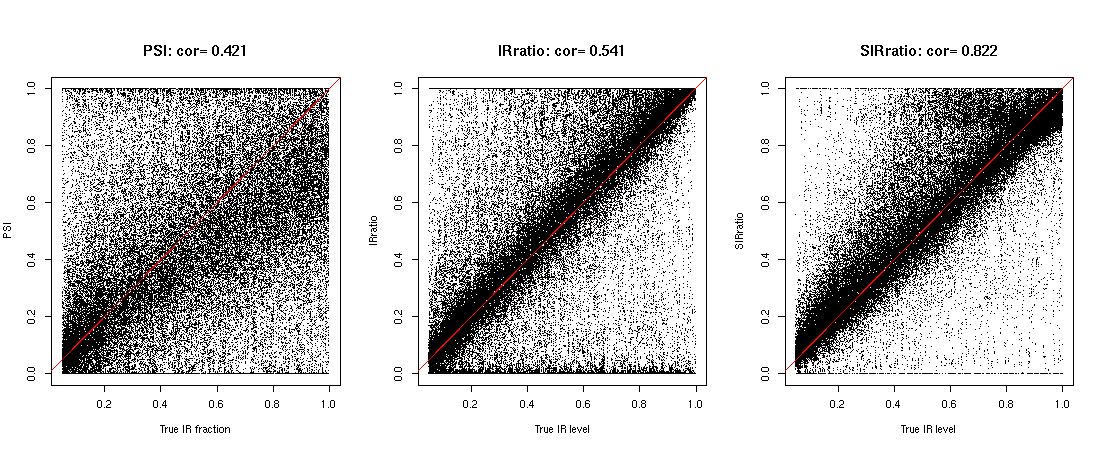

**Supplementary Table 1: Summary statistics of distances to the true value for over all simulated datasets.** True IR-levels and estimates from the three methods were ilr-transformed. For each retained intron, the distance was then calculated as the absolute value:

distance to the true value = |ilr( true value ) - ilr( estimate )|

|  | **Distance to the ground truth**  **(Pooled simulated data)** | | | | |
| --- | --- | --- | --- | --- | --- |
| **Method** | **1st quantile** | **Median** | **Mean** | **3rd quantile** | **sd** |
| **PSI** | 0.142 | 0.332 | 0.811 | 0.714 | 2.22 |
| **IRratio** | 0.064 | 0.184 | 0.609 | 0.867 | 0.81 |
| **SIRratio** | 0.050 | 0.119 | 0.250 | 0.279 | 0.38 |

**Supplementary Figure 2: Violin plots of differences to the true value for each scenario (Simulated data).** The three IR measures were computed under five IR-level bins ([0.05-0.15], [0.2-0.4], [0.4-0,6], [0,6-0,8], [0.8-1]) and three gene coverage bins (LOW, MEDIUM, HIGH), thus giving 15 scenarii. Each retained intron is evaluated exactly once under each scenario. All IR-levels were ilr-transformed. For each retained intron, the difference was obtained by:

difference to the true value = ilr( true value ) - ilr( estimate )

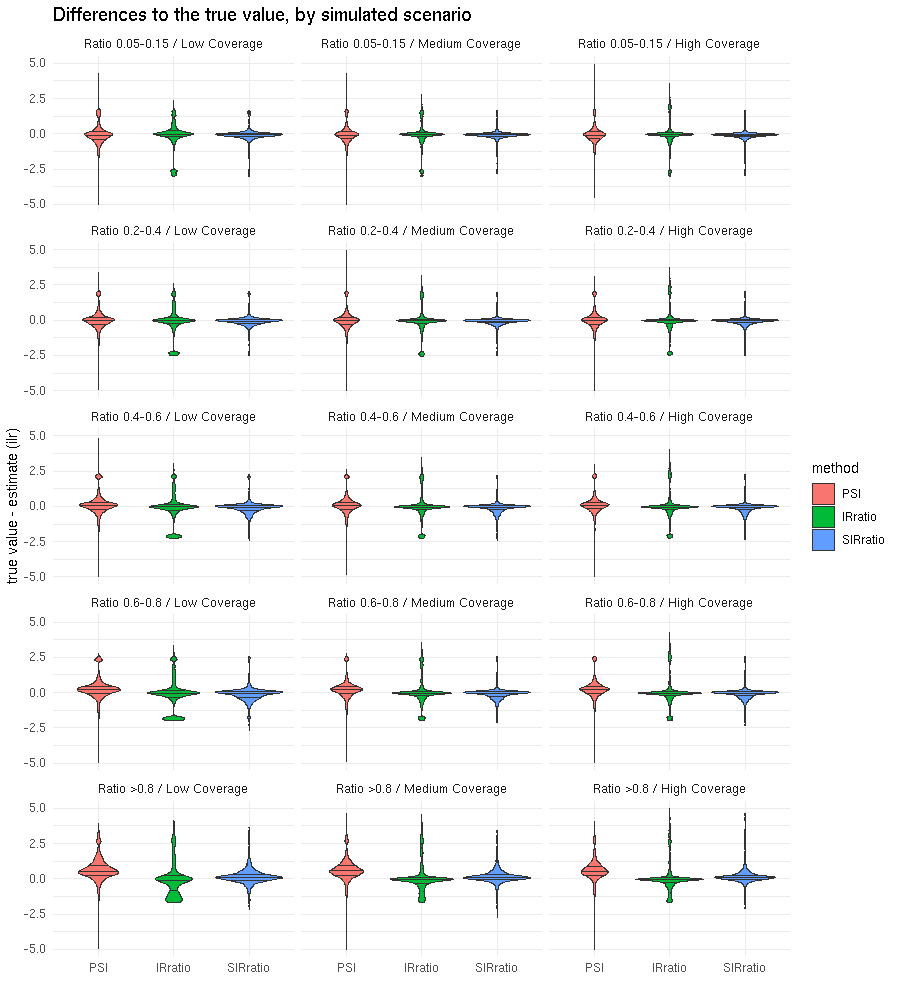

**Supplementary Table 2: Overall distances between IR-level estimates between the technical replicates (Real data).** Following [[9]](https://www.zotero.org/google-docs/?ozDNrM), we quantified reproducibility through the distances between IR-levels obtained in each replicate. Distances were computed on ilr-transformed ratios across the five technical replicates for each independent intron identified from the reference transcriptome.

|  | **Overall distances between technical replicates** | | | | |
| --- | --- | --- | --- | --- | --- |
| **Method** | **1st quantile** | **Median** | **Mean** | **3rd quantile** | **sd** |
| **PSI** | 0.138 | 0.352 | 2.932 | 0.920 | 6.17 |
| **IRratio** | 0.076 | 0.173 | 0.245 | 0.319 | 0.30 |
| **SIRratio** | 0.057 | 0.146 | 0.229 | 0.283 | 0.34 |

**Supplementary Figure 3: Pairwise boxplots of distances between IR-level estimates between the technical replicates (Real data).**

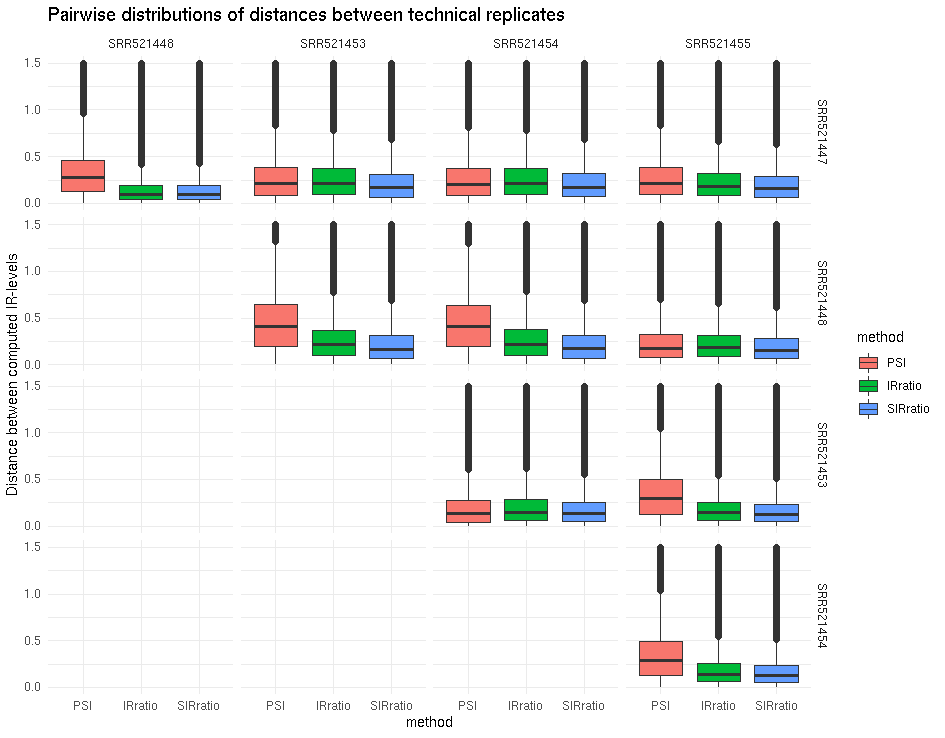

**Supplementary Figure 4: Global Pearson and Spearman correlation between technical replicates (Real data).**

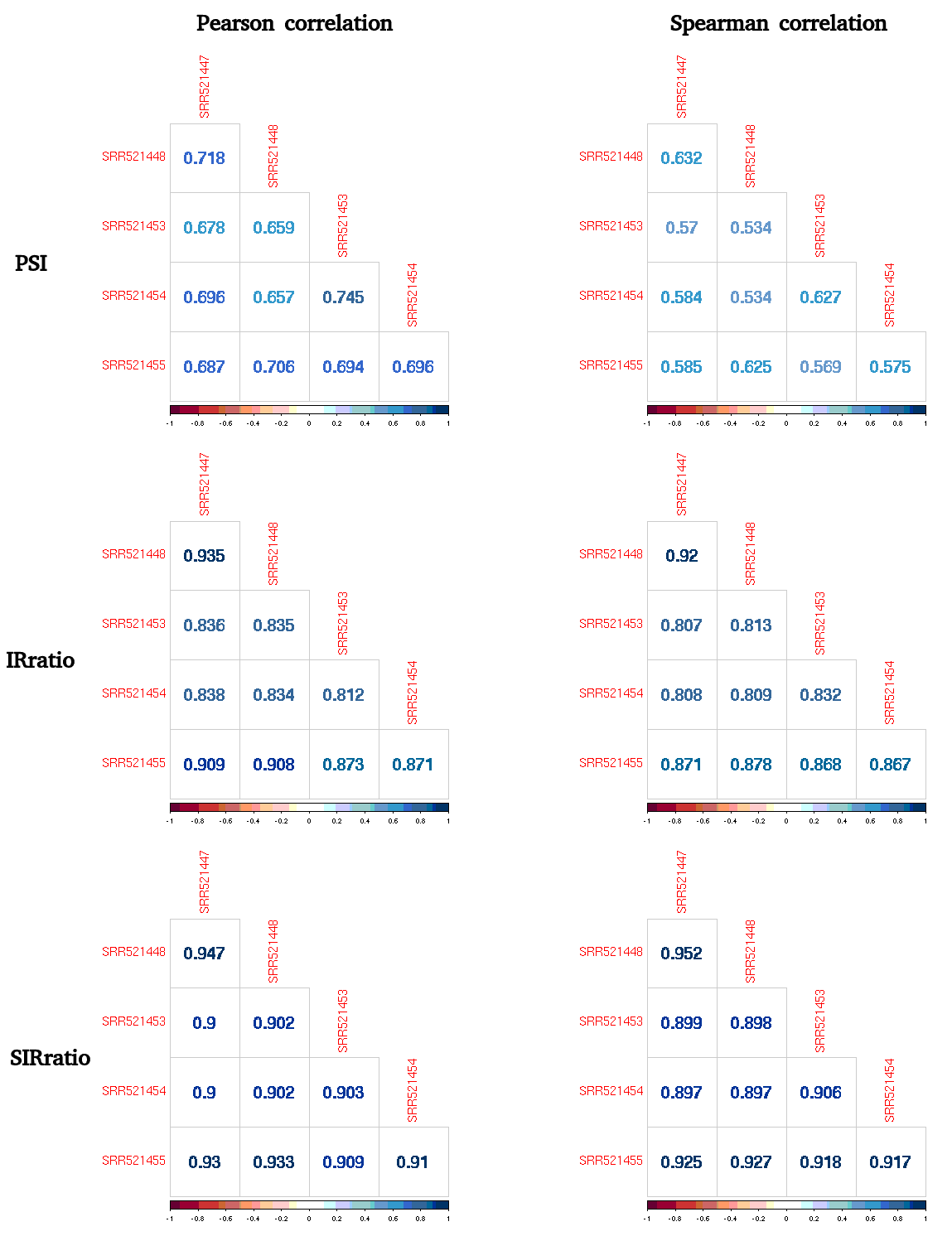

**Supplementary Table 3: Distances between estimates and reference values computed from the matched long read sequencing experiment.** We used ratios computed from long read data as reference values**.** Euclidean distances between reference and evaluated IR measures were calculated on ilr-transformed ratios. That is, for each intron:

distance to reference = |ilr(reference value) - ilr(IR measure)| .

|  | **Median distance to the reference value** | | | | | |
| --- | --- | --- | --- | --- | --- | --- |
| **Method** | **SRR521447** | **SRR521448** | **SRR521453** | **SRR521454** | **SRR521455** | **Pooled** |
| **PSI** | 0.479 | 0.415 | 0.887 | 0.944 | 0.418 | 0.387 |
| **IRratio** | 0.272 | 0.254 | 0.344 | 0.322 | 0.296 | 0.275 |
| **SIRratio** | 0.224 | 0.202 | 0.259 | 0.251 | 0.226 | 0.214 |

**Supplementary Figure 7: IR-levels computed with IRFinder on the EMT time course dataset. A.** Principal component analysis of IRratio values across time points**. B.** Heatmap of IRratio values and hierarchical clustering on the 100 most influential retained introns on the first component, selected by sparse PCA. Due to unstable estimates of IR-levels creating additional unwanted variations, classical methods for analyzing high-dimensional data struggle to cluster biological replicates together and select relevant intron biomarkers of EMT stages.

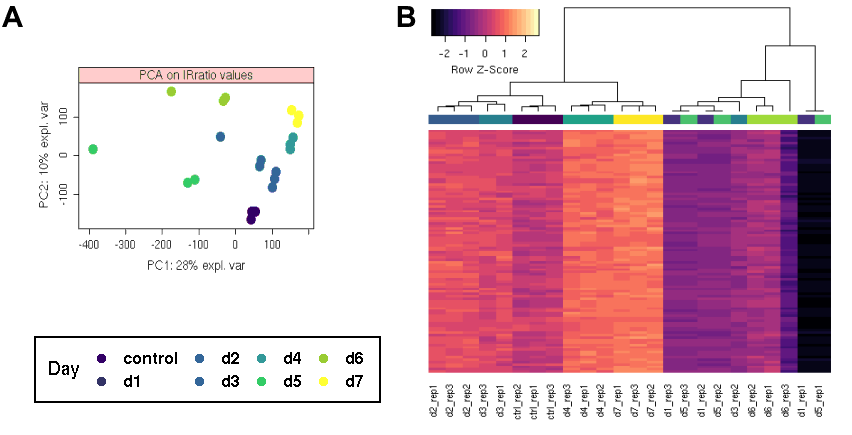

**Datasets used for evaluating IR-level estimates**

For all computations and analyses, we made use of the reference genome hg38 and of the reference transcriptome *Homo sapiens* (release 97) from the ENSEMBL database.

**Simulated Data**

To investigate the properties and performances of the three IR measures, we computationally generated an RNA-seq experiment with known IR rates, and varying flanking exon skipping and gene coverage.

**Gene and transcripts selection.** For each annotated and multi-exonic gene of the human chromosome 1 (1566 loci), we randomly selected one of its multi-exonic transcripts (called *canonical transcript*). Then an intron was selected randomly in this transcript to define an *IR-transcript* for the gene (cf: **Supplementary Figure 8-A**).

**Gene coverage and transcript usage.** To allow comparison across gene expression levels (LOW=0-50 reads per base, MEDIUM = 50-250 reads per base,, HIGH=250-1500 reads per base) and retention rates ([0.05-0.15], [0.2-0.4], [0.4-0,6], [0,6-0,8], [0.8-1]), we simulated 15 conditions with 3 replicates each, so that each intron was evaluated under each of the 3 GENE x 5 IR settings (cf: **Supplementary Figure 8-B**). Actual values for IR rates and gene coverage depth were selected uniformly at random in each bin range.

**Read simulation.** Short RNA-seq reads were simulated using the R package Polyester [[10]](https://www.zotero.org/google-docs/?40XRZN). All R scripts designed for generating the simulated datasets are made available at: <https://github.com/lbroseus/SimIR/>.

**Supplementary Figure 8: Illustration of the simulated RNA-seq experiment. A**) For each multi-exonic gene on the human chromosome 1 (1 566 loci), we drew one of its (multi-exonic) transcripts (*canonical transcript*). Then an intron was randomly selected as well as one of its flanking exons, to define an *IR-transcript* and a transcript with alternative exon skipping. These canonical, IR and exon-skipping transcripts constitute the exact transcriptome expressed in our simulated RNA-seq experiment. **B**) To detect whether the performances of the three IR-measures would vary depending on read depth or on the true IR-levels, we considered and simulated 15 “Coverage x IR scenarii”. The quantification of each selected IR-transcript was evaluated under all 15 scenarii.

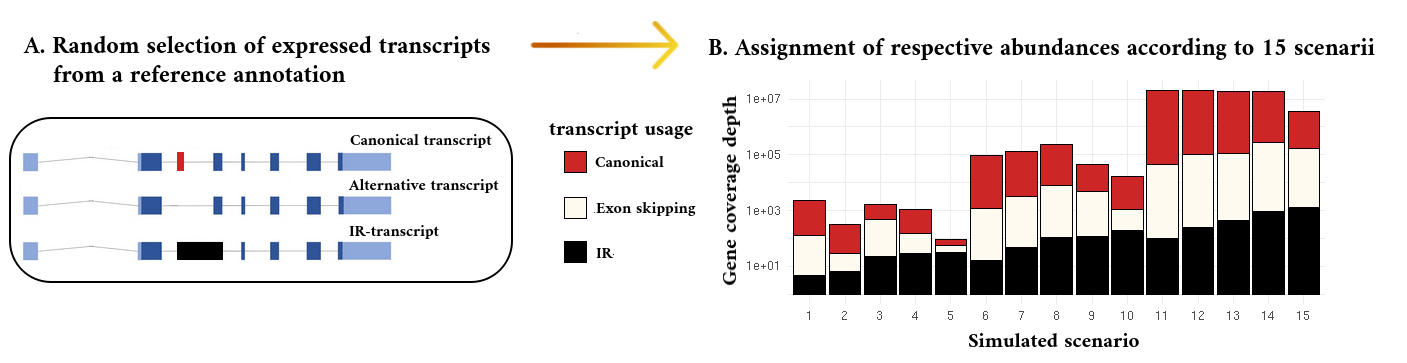

**Real Data**

We downloaded all 10 technical replicates from a Poly-A+ RNA-sequencing paired-end experiment ([SRX159821](https://www.ncbi.nlm.nih.gov/sra/SRX159821%5Baccn%5D)) of the GM12878 cell line [[11]](https://www.zotero.org/google-docs/?abmukG). Data quality was checked using FastQC, which led to the exclusion of five runs (SRR521449, SRR521450, SRR521451, SRR521452, SRR521456) due to bad quality warnings. The five remaining runs we kept for analysis are listed in **Supplementary Table 4** below. RNA-seq samples were trimmed for adapters using the dedicated Perl script provided with IRFinder.

**Supplementary Table 4: Real Illumina RNA-seq datasets used to evaluate IR measures.** We report alignment rates obtained with Bowtie2 with parameters suggested by KMA.

| **SRA id** | **Library Size**  **(# of reads after trimming)** | **Alignment rate**  **(Bowtie2)** |
| --- | --- | --- |
| SRR521447 | 23,855,992 | 96.91% |
| SRR521448 | 27,498,369 | 97.13% |
| SRR521453 | 8,313,056 | 95.74% |
| SRR521454 | 8,316,090 | 95.89% |
| SRR521455 | 37,187,942 | 96.82% |
| Pooled experiment | 105,171,449 | 96.77% |

**Validation using real Oxford Nanopore Long Read.** We downloaded the whole NA12878 RNA-direct Oxford Nanopore experiment made available by the Nanopore Consortium (<https://github.com/nanopore-wgs-consortium/NA12878>). It contains over 10 million raw sequences generated from the human cell line GM12878 [[12]](https://www.zotero.org/google-docs/?qvAPXg). Long reads were aligned to the human reference genome (hg38) using Minimap2 [[13]](https://www.zotero.org/google-docs/?S7EpKE).

Reference IR-levels were calculated from the long read dataset as follows. For each independent intron derived from the annotation, we recorded the number of long reads spanning the intron (*overall read abundance*) as well as the number of long reads containing the intron (*intronic read abundance*). IR levels were then obtained as the proportion: intronic abundance / overall abundance. As low counts may lead to unreliable estimates of proportions [[14]](https://www.zotero.org/google-docs/?XsWj5e), we required that overall read abundance ≥ 30, intronic read abundance ≥ 20 and a ratio ≥ 0.05 to include the intron in the analysis. All in all, the comparison with long reads was made over 703 annotated introns. All R functions used to compute IR-levels from long read data are implemented in the R package S-IRFindeR.

**Supplementary Figure 9: Reference IR-levels computed from the NA12878 direct-RNA Oxford Nanopore experiment.** In total 703 reference IR-level values were obtained from long reads (LR) and used for comparison to the measures derived from short read RNA-seq in the same cell line.

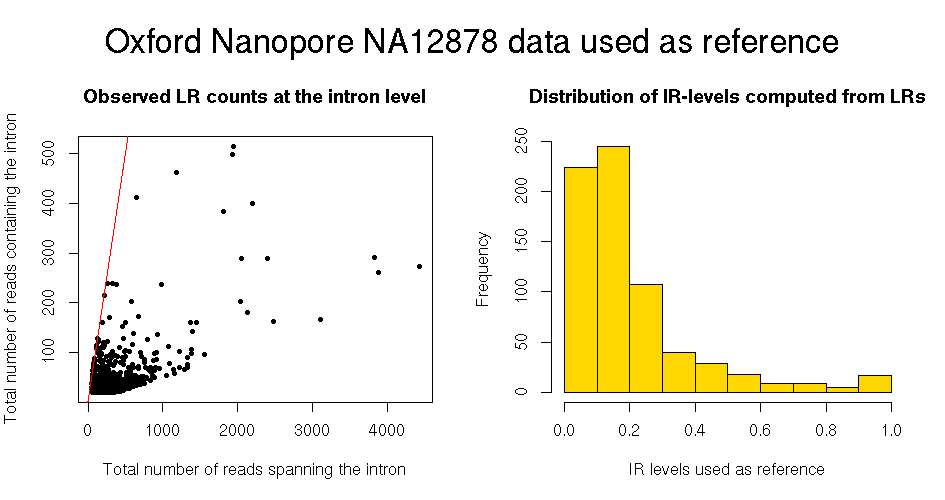

**Application of SIRFindeR to a time series RNA-seq experiment on the human cell line H358.** We downloaded the 24 Poly(A) RNA-seq samples from the H358 cell lines generated in the study [[15]](https://www.zotero.org/google-docs/?r0SlDc). Cells were treated with doxycycline to induce epithelial-to-mesenchymal transition (EMT) and followed during 7 days. The RNA-seq experiment is a time series of triplicates divided into 7 time points (days) and a control (untreated). The data are paired-end reads sequenced on Illumina Hiseq 2000 or Hiseq 2500.

**Methods**

**Refining intron annotation using RNA-seq data**

We first derive known *intron intervals* from a reference annotation. All annotated overlapping exons are merged, and introns are defined as the interspersed genomic intervals that overlap with no merged exons. In addition, potential alternative «*de novo*» intron borders are detected from the RNA-seq dataset using STAR [[16]](https://www.zotero.org/google-docs/?rEizs0) alignments (**cf: Supplementary Figure 10**). By default, we consider only “novel”exon-exon junctions supported by at least 5 spliced reads, defining an intron of at least 30 bp. These parameters can be adjusted by the user.

We then combine both annotated and *de novo* introns:

1. we crop the genomic coordinates of annotated introns having supported alternative 3’ or 5’ donor sites to obtain a list of *data-curated independent introns;*

*2.* novel introns that fall entirely into an exon can also be added to the analysis.

**Supplementary Figure 10: Illustration of S-IRFindeR’s approach to polish genomic coordinates of annotated introns.** In this example, three transcript isoforms are expressed but only the first transcript is present in the reference annotation. The alternative exon-exon junction from the second transcript can be detected from read alignments (spliced reads) and used to update the genomic coordinates of the intron (brown arrow).

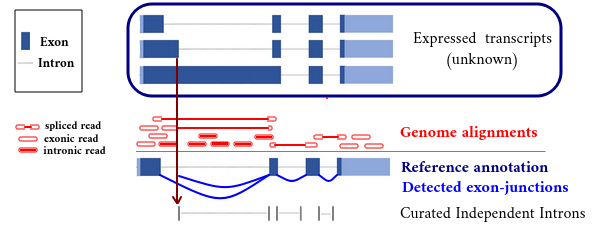

**The SIRratio**

There are two types of directly informative reads for IR: reads spliced across the intron [[17,4,18]](https://www.zotero.org/google-docs/?4rua57), which originate from transcripts without the intron, and reads overlapping the intron. The IRratio uses only informative reads to measure the proportion of IR-transcripts.

Precisely: $\frac{MedianIntronReadDepth}{MedianIntronReadDepth+SpliceCount}$.

where *SpliceCount* is the number of reads spliced across the intron, and is taken as an estimate of the depth of transcripts that do not contain the intron.

**Supplementary Figure 11. Comparison between true splicing events and intronic abundances and their estimates by IRFinder (Simulated data).** Global spearman correlation coefficients are indicated. The left-hand plot, confronting the number of spliced reads (SpliceCount) to the true abundance of spliced transcripts, highlights SpliceCount’s tendency to under-estimate the fraction of spliced transcripts. The right-hand plot shows how well the median intronic depth tallies to the true intronic abundance.

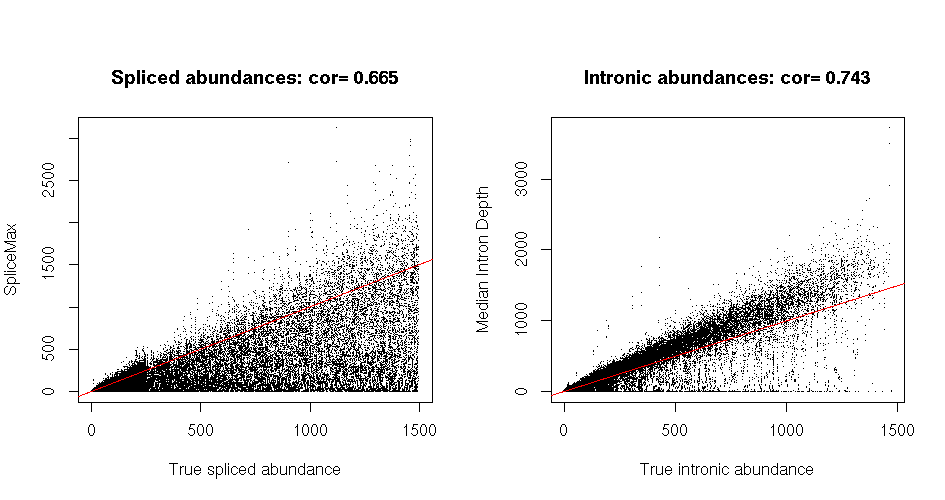

The SIRratio, similarly to the IRratio, evaluates intronic and normal splicing abundances using only spliced reads and intron-overlapping reads [[4,18]](https://www.zotero.org/google-docs/?5eIF2V). However these measurements are adjusted using a shrinkage and resampling approach.

**Estimating the abundance of spliced transcripts.** Using our simulated experiment, where the true transcript fractions are known, we noticed that SpliceCount values are overdispersed and tend to underestimate the true value (cf: **Supplementary Figure 11)**.

As a means to control for overdispersion, we choose to regularize SpliceCount values using a *shrinkage* approach [[19,20]](https://www.zotero.org/google-docs/?uaLD4r). More precisely, for any intron from the gene, we will estimate its spliced abundance as a compromise between the point estimate SpliceCount and the overall gene coverage depth (GeneDepth), that is: splicing abundance = (1-λ)*SpliceCount + λ*GeneDepth, where λ is the degree of shrinkage towards the estimate of the gene coverage depth (between 0 and 1).

**Estimating gene coverage depth**. We treat each gene separately, and consider all its annotated and *de novo* exon-intron junction sites. At each site, we compute the number of spliced reads (SpliceCount) and the number of overlapping reads (ExonToIntronCount). The non-zero sums SpliceCount+ ExonToIntronCount provide as many point estimates for the gene coverage depth. We then make use of the resampling technique called Bootstrap, to integrate these values into a more stable estimate for GeneDepth [[21]](https://www.zotero.org/google-docs/?PPnMG9). Specifically, we create 100 *bootstrap samples* of size 50 by drawing with repetition in the set of point estimates. The 100 mean values computed from these bootstrap samples constitute a *bootstrap distribution*, whose mean, ${\mu_{boot}}$, is taken to estimate GeneDepth.

**Degree of shrinkage**. The parameter λ is calculated automatically for each intron. In a sense, it measures the degree of confidence we have in ${\mu_{boot}}$for reflecting the abundance of splice transcript better than the sole SpliceCount value. Our rationale for computing λ is the following. The more informative (ie: non-zero count values) exon-intron count values the better the estimate of GeneDepth. This is accounted for by putting a weight $\frac{n}{n+1}$on GeneDepth. Secondly, the closer the gene coverage depth to the intronic depth, the worst it estimates the abundance of spliced transcripts. For this reason, we apply the penalty: ${\mu_{boot}}-intronicdepth\vee\frac{}{{\mu_{boot}}+intronicdepth}$.

Eventually, we take: λ = $\frac{n}{n+1}$x ${\mu_{boot}}-intronicdepth\vee\frac{}{{\mu_{boot}}+intronicdepth}$,

where *n* is the number of informative exon-intron junctions for the gene.

**Estimating intronic abundances.** The IRratio relies on the median coverage depth of introns to quantify intron-retaining transcripts. However, we noticed that median values can be unstable, displaying drastic differences between technical replicates, and give aberrant estimates of intronic abundances (eg: zero values whereas many reads map within the intron, cf: **Supplementary Figure 11)**. For this reason, we choose to consider instead the number of reads overlapping the intron (*intronic count*) that we integrate in a formula which takes intron width into account.

**SIRratio formula.** For a given intron, let: ꭤ = Integer( $\frac{intronwidth}{readlength}$) (the integer part)

and Ⲭ = $\frac{introniccount}{introniccount+splicingabundance}$

Then, the SIRratio is computed as: SIRratio = $\frac{Ⲭ}{ꭤ+1-ꭤⲬ}$.
